# Interpreting pathogen genetic diversity during measles epidemics

**DOI:** 10.1101/2020.01.30.926998

**Authors:** CJ Worby, BA Bozick, PA Gastañaduy, Luojun Yang, PA Rota, BT Grenfell

## Abstract

While measles remains endemic in parts of the world, efforts to eliminate measles transmission continue, and viral sequence data may shed light on progress towards these goals. Genetic diversity has been used as a proxy for disease prevalence; however, seasonally-driven disease dynamics are typically characterized by deep population bottlenecks between epidemics, which severely disrupt the genetic signal. Here, we simulate measles metapopulation dynamics, and show that it is the population bottleneck, rather than epidemic size, which plays the largest role in observed pathogen diversity. While high levels of vaccination greatly reduces measles diversity, paradoxically, diversity increases with intermediate levels of vaccination, despite reducing incidence. We examined diversity and incidence using published data to compare our simulated outcomes with real observations, finding a significant relationship between harmonic mean incidence and genetic diversity. Our study demonstrates that caution should be taken when interpreting pathogen diversity, particularly for short-term, local dynamics.

## Introduction

Measles elimination remains a key public health goal for many regions around the world (1). Nevertheless, it remains endemic in many countries including China, India and many countries in sub-Saharan Africa, and sporadic reintroductions to countries which have previously eliminated the disease have resulted in large outbreaks. Measuring progress towards local disease elimination is an essential component of coordinating optimal vaccination efforts both regionally and globally. However, surveillance is often incomplete in regions with fewer resources to implement widespread vaccination, and as such, the magnitude of disease circulation may be underestimated. Furthermore, it can be challenging to determine whether a given set of cases represents a continuation of endemic transmission chains, or a novel strain imported from another region. Genomic surveillance has been promoted in recent years as a tool which may be used to provide insight into unobserved disease transmission dynamics, through identification of local transmission clusters and discrimination between imported and endemic cases. This information can help measure the impact of changes to vaccination policy. In this study, we explore the relationship between pathogen genomic diversity and epidemiological dynamics for a seasonally driven disease such as measles.

In addition to the public health importance of reducing the global morbidity and mortality caused by the pathogen, there are several reasons why we have chosen to explore diversity and disease dynamics within the context of measles transmission. Firstly, the epidemiological dynamics of measles have been extensively studied, and can be modeled within a well-understood framework (2-5). Secondly, measles virus is not thought to have complex evolutionary dynamics. Immunity from both infection and vaccination is long-lasting and protects against all variants (6, 7); as such, selective pressure from adaptive immunity is thought to be minimal (8). Finally, measles dynamics vary considerably across different scenarios, encompassing biennial and annual outbreaks, as well as chaotic dynamics (9), providing a range of settings to explore how genetic diversity is impacted by population fluctuations.

Measles virus has just a single serotype, although 24 genotypes have been defined (10) on the basis of (at minimum) the 450-nucleotides coding for the COOH terminal 150 amino acids of the nucleoprotein (N-450) (11). The World Health Organization (WHO) Global Measles and Rubella Laboratory Network supports measles genotyping on a global scale. At present, laboratory confirmation of measles infection is based on detection of virus specific IgM antibodies in serum samples. Though WHO encourages the collection of specimens for virus detection, the quality and completeness of virologic surveillance varies considerably by country and WHO region (12). Classification of strains sampled during an outbreak into these genotypes can provide additional insights into epidemiological dynamics. In many countries with high vaccine coverage, a relatively high turnover of predominant genotypes is observed, since outbreaks are generated by sporadic introductions, and are subsequently either eliminated, or reduced to a low level, allowing new importations of different genotypes to take over (13). In contrast, countries with lower vaccination coverage and endemic measles are more likely to be characterized by a single genotype (14). For instance, measles in China is predominantly caused by genotype H1, with only few importation-associated cases identified belonging to other genotypes (15, 16), while genotype D8 dominates in India (17). Changes in genotype distribution can be indicative of breaks in transmission and importation (13), and new outbreaks in a previously disease-free population are likely to comprise a single major genotype and exhibit little genetic diversity (18). However, genotype replacement may also occur by chance with limited or no vaccination coverage (19).

While genotype-level data can provide important insights into broad population dynamics, it is hoped that whole genome sequence data may provide finer details, particularly within endemic populations dominated by a single genotype. Phylodynamic analyses can identify patterns in population dynamics based on the inferred evolutionary history of sampled lineages (20). Tools such as BEAST2 (21) allow the reconstruction of demographic history using sequence data, and can successfully identify fluctuations in effective population size in a variety of scenarios. However, populations which experience repeated population contractions, or bottlenecks (such as seasonally-driven disease outbreaks), will periodically lose a large proportion of lineages, and the genetic signal from one period may be largely uninformative about the previous period (8).

Previous studies have suggested that declining genomic diversity is indicative of progress towards measles elimination and/or success of additional vaccination efforts (13, 22, 23). However, linking observed patterns in genetic diversity to population dynamics can be challenging, since a multitude of factors, including geographic and temporal sampling bias, selection pressure and interaction with external populations, can impact observations. It has been noted that while levels of genetic diversity and vaccination coverage may be linked, the factors promoting viral diversity are not well understood (24). Furthermore, stochastic effects can have a key role in genotype replacement (19), and may also play a role in fluctuations of within-genotype genetic diversity.

Genetic diversity is proportional to the expected time to most recent common ancestor and the mutation rate of the organism (25). At equilibrium, nucleotide diversity approaches a level of 2N_e_*µ* in a haploid population of fixed size, undergoing neutral evolution (where N_e_ is the effective population size, and *µ* is the per-generation mutation rate). Within a monotonically increasing or decreasing population, the genetic diversity is expected to change proportionally to this size. However, the evolutionary dynamics expected within a temporally fluctuating population are more complex. If the population size is not constant, then the diversity is strongly dictated by the size of recent population minima, or bottlenecks. The harmonic mean captures this, and is approximately equivalent to the effective population size over time (26). While the genetic diversity will increase as the population expands during a large outbreak, much of this diversity may be lost in the subsequent trough. As such, the interpretation of genetic data with respect to both long-term and short-term periodic population dynamics is challenging. We hypothesize that the pathogen genetic diversity observed from a population is governed by the harmonic mean incidence, rather than either peak or cumulative incidence.

In this study, we aimed to explore the relationship between measles population dynamics and observed levels of genetic diversity within a population. Stack et al. demonstrated the loss of genetic signal between epidemics, and the importance of sampling strategy in recapturing temporal dynamics of measles (8). We built upon this framework, initially exploring genetic diversity under pre-vaccination epidemic dynamics, verifying whether the relationship between harmonic mean population size and diversity first described by Nei (26) holds in this context. We then investigated the impact of vaccination introduction, and considered a meta-population model with varying levels of mixing. Simulation of evolutionary and epidemiological dynamics allowed us to establish whether a clear relationship between neutral viral evolution and epidemiological dynamics could be identified. Finally, we determined the extent to which real data follow the patterns we expect in theory using measles sequence data collected from GenBank.

## Methods

### Epidemic Model

The framework of our epidemic and evolutionary model is similar to that presented by Stack et al. (8). We simulated measles epidemic dynamics using a seasonally forced SIR model with homogeneous mixing (2). This is a stochastic, discrete-time model with two-week time steps, reflecting the measles generation time. We first derived model parameters fitted to data from pre-vaccine era London, using the tsiR package (27) in R to provide baseline population dynamics, resulting in large, biennial epidemics. For each new infection, we drew a source of infection at random from the pool of infectives in the previous generation, storing this information in order to generate evolutionary dynamics along paths of transmission (8).

### Evolutionary Model

Evolutionary dynamics were simulated under a neutral model. Since we are interested in population-level, rather than individual-level, dynamics, we do not consider within-host diversity, characterizing each host by a single genotype which may accumulate new mutations upon transmission (eg. (8, 28)). We generate sequence data for 0.1% of the total number of cases, randomly selected across a given time period. We found that this sampling fraction was sufficient to capture true genetic diversity, and required significantly less computational time than larger sampling proportions. We experimented with both proportional and constant sampling rates. For N samples, we can generate a minimal spanning transmission tree, such that for any given pair of hosts, we can trace their ancestry back to a most recent common ancestor. To generate sequence data for the N samples, we first identified the set of common ancestors, or coalescent points on the transmission tree, resulting in at most N(N-1)/2 hosts. To generate a sequence at a given node, we introduced *m*∼Pois(*µk*) polymorphisms relative to its preceding coalescent point, where *µ* is the expected number of mutations between infector and infectee, and *k* is the length of the transmission chain between the two nodes. We simulated a burn-in period of 2000 time steps (∼77 years) prior to sampling genomes, which in all scenarios was sufficient to contain the most recent common ancestor of sampled hosts. We calculated sampled nucleotide diversity across time to evaluate the evolutionary dynamics in the population.

We calculate the harmonic mean of a population as a sliding window of length *k*, i.e.

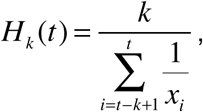

where *x*_*i*_ is the population at time *i.* According to population genetics theory, this value should be proportional to the expected genetic diversity within the population (26), and we aimed to investigate to what extent this holds under various epidemiological scenarios. We further considered the time to most recent common ancestor (t_MRCA_) of all samples at a given time as more general metric to measure a population’s capacity for genetic diversity.

### Vaccination

We considered the effect of vaccination on the neutral evolutionary dynamics of the pathogen in a single population of size 3,000,000 by scaling the relative birth rate ρ (rate at which new susceptible individuals are introduced to the population) such that *v*_*t*_=1-ρ, where *v*_*t*_ is the vaccination coverage at time *t*. We used a simple model assuming that the vaccine provides complete and long-lasting immunity against all strains (6), and that vaccine escape does not occur (7). We considered both immediate rollout of vaccination, whereby from time t_vacc_ onwards, 50% of the population are considered vaccinated, as well as scaling up of vaccination, in which the rate of vaccination linearly increases from 20% at t_vacc_ to 50% 10 years later. In order to explore general dynamics, we did not allow the population to go extinct, instead generating at least one new case in each generation. Pathogen extinction is likely to occur when a closed population is below the critical community size (29). For measles, the critical community size is 250,000-400,000 (30), and in practice, we found that extinction was rare in our large population while vaccination coverage was <90%.

### Metapopulation Model

We next considered the evolutionary dynamics of measles in a multi-population setting, in which we model dynamics in *x* cities with population movement. Initially we considered equally-sized population pairs, experiencing either in phase or out of phase biennial epidemics. We then considered introducing vaccination into one population to determine the effect on the neighboring city, under a range of vaccination coverages and inter-population movement rates. The rate of movement of infected hosts from A to B is proportional to the disease prevalence in A, and we define *θ* to be the expected number of infected hosts to migrate during peak incidence.

More generally, we a used a core-satellite metapopulation model (4), with one major city, population *n*_*1*_=3,000,000, and *x*-1 neighbouring cities with smaller populations *n*_*2*_, …, *n*_*x*_, ranging from 0.5% and 60% the size of the major city. We assumed that the cities were equidistant from one another, and that movement between any two cities was driven by their population sizes alone. We calibrated the rate of population movement such that the pre-vaccination critical community size approximately matched historic records (3). In the metapopulation setting, we did not prevent local populations from going extinct, since they may be reseeded from neighboring populations.

### Genbank sequence data

To explore the relationship between observed genetic diversity and regional incidence, we acquired all publicly available measles sequence data from GenBank (31), and compared this to national surveillance data. To obtain the sequence data, we used the search term “nucleoprotein” and filtered by organism “measles morbillivirus”, and sequence length greater than or equal to 450 in order to acquire (at least) the standard N-450 gene sequence (32). We further removed entries lacking geographic or temporal metadata, leaving 9169 strains. We used MUSCLE (33) to perform multiple sequence alignment, and the R package ‘ape’ (34) to calculate genetic distances post-alignment. We used monthly measles reporting data from China (35), India (36), Europe (37) and the USA (38), allowing us to compare the annual sampled genetic diversity with both general trends in incidence and the harmonic mean (with a two-year sliding window).

## Results

We first considered a single population, with the characteristics of pre-vaccine era London, to consider evolutionary dynamics in the absence of importation and intervention. This population experienced large, biennial epidemics with deep inter-seasonal troughs. Figure 1 shows the incidence and sampled time to most recent common ancestor (t_MRCA_) under various vaccine coverage levels. The time to most recent common ancestor is proportional to the expected diversity within a population, and the fluctuations corresponding to biennial outbreaks are apparent under the baseline scenario (Figure 1A). However, while average t_MRCA_ exhibits clear periodic dynamics, considerable variance exists and any given simulation may exhibit large fluctuations as a result of the randomly selected lineages which persist through a bottleneck.

**Figure 1.**
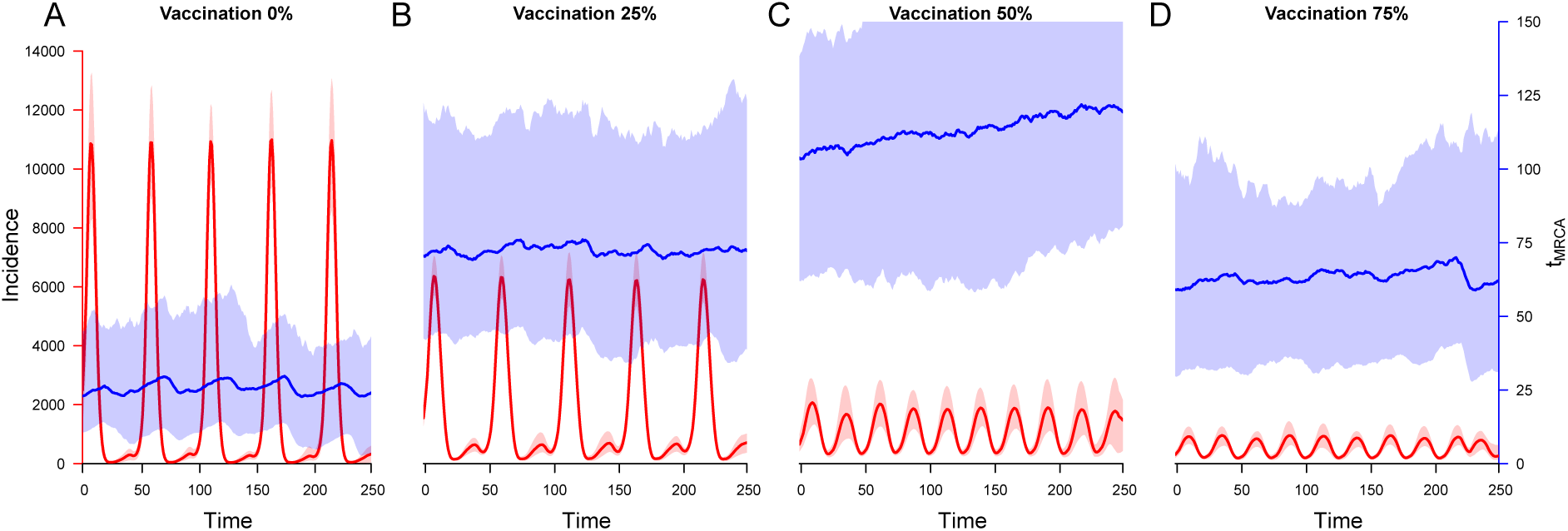
Simulated measles incidence (red) and mean time to most recent common ancestor (t_MRCA_) (blue) in a single population. We varied vaccination coverage and plotted the average incidence and t_MRCA_ over a period of ten years, averaging both across 50 simulations. Shaded areas represent the interquartile range.

### Impact of vaccination

Figures 1B-D show the effect of increasing vaccination coverage on the epidemic dynamics and t_MRCA_. The expected t_MRCA_, and therefore genetic diversity, increases for intermediate levels of vaccination, but falls as vaccination coverage exceeds 50%. The periodicity of t_MRCA_ observed under the baseline scenario is lost as vaccination levels increase, and variation also increases.

The timing of vaccination introduction affected short-term dynamics, as expected from previous findings (39). Providing immediate 50% coverage at peak incidence resulted in a severe post-epidemic bottleneck, and a high probability of extinction. In contrast introduction between epidemics was much less likely to result in extinction (Figure S1A). While the short-term effects on the population differed, providing the population did not reach extinction, the long-term dynamics were similar. We found that the post-vaccination population harmonic mean was larger under 50% vaccination coverage, allowing for the accumulation of greater genetic diversity. Timing was less important for gradual introduction of vaccination, and the long-term dynamics were equivalent to those following immediate introduction (Figure S1B).

More generally, we considered varying ρ between zero (no new susceptibles introduced; total vaccine coverage) and two (twice the baseline rate of susceptible introduction, equating to an increased birth rate), while also simulating evolutionary dynamics. Figure 2 shows the impact of altering this rate on the genetic diversity and the epidemic dynamics. Epidemics shift from biennial to annual as vaccination coverage is increased (Figure 1, 2A), though the population bottleneck increases for coverage up to 50%, as reflected by the average t_MRCA_ (Figures 1A-D) and the population harmonic mean (Figure 2B). With larger population bottlenecks, the periodic fluctuations in population are not reflected in the average t_MRCA_ (Figure 1). We calculated the mean genetic diversity across the simulated population, finding it to be well correlated with the harmonic mean (Figure 2C), as anticipated according to theory (26). As such, we found that genetic diversity was expected to increase as vaccination coverage increased up to around 50%, before reducing for higher levels of coverage (Figure 2D). Moderate increases in birth rate led to more severe bottlenecks between epidemics and a lower genetic diversity, although a greater probability of extinction, while larger increases shifted the epidemic dynamics from biennial to annual, increasing the bottleneck size.

**Figure 2.**
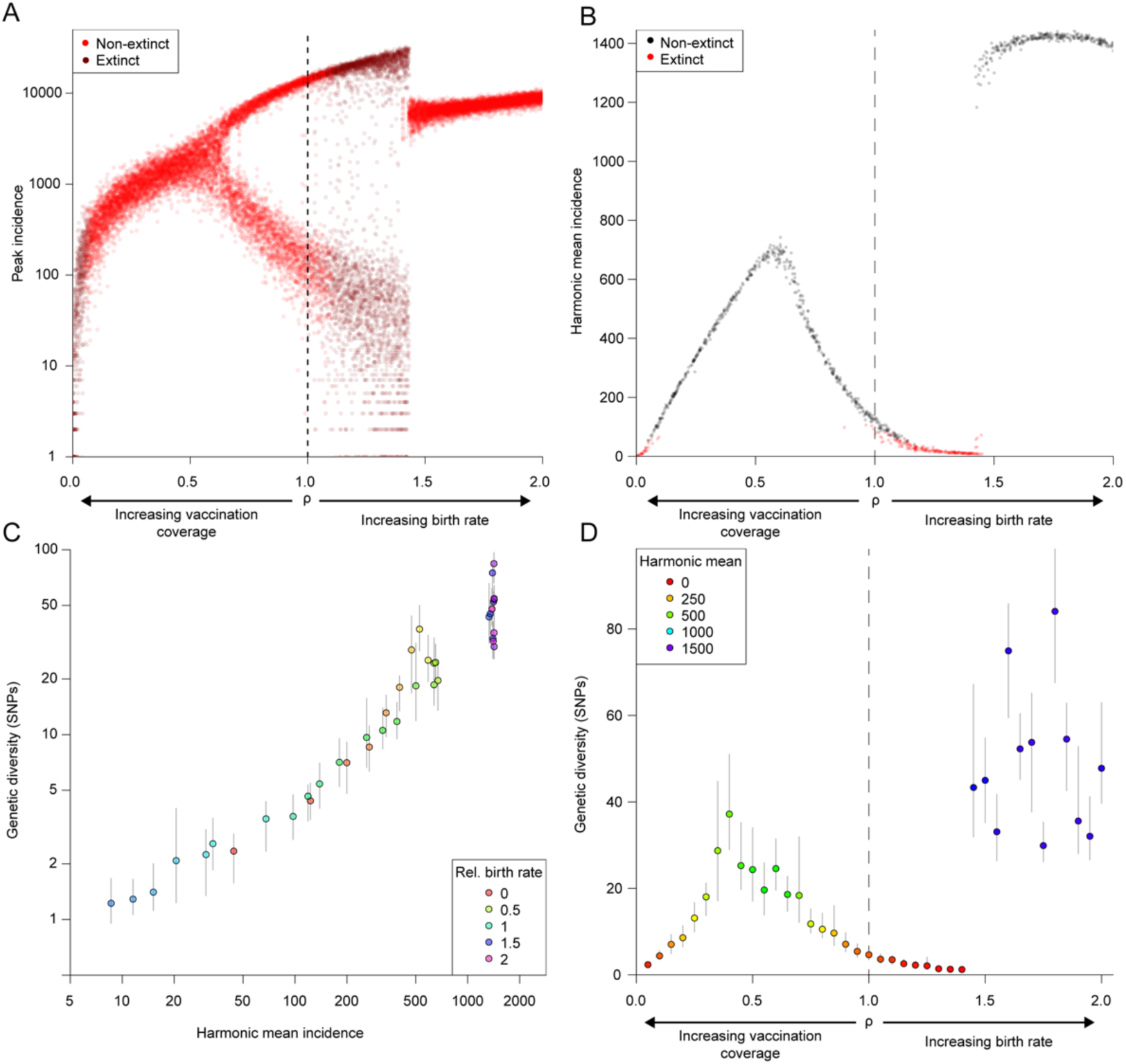
Vaccination affects harmonic mean incidence and expected genetic diversity. Vaccination levels were varied relative to the baseline to explore the effects on population and evolutionary dynamics. Simulations for which incidence reaches 0 are reseeded, but are recorded as having gone extinct. (A) Bifurcation diagram illustrating the annual peak incidence for varying ρ. (B) The harmonic mean (across a simulation period of 1000 biweeks) for varying ρ. Simulations which would have gone extinct without artificial propagation are marked in red. (C) Correlation between simulated genetic diversity and harmonic mean. (D) Simulated genetic diversity for varying levels of vaccination.

While our baseline transmission rate parameters were derived from pre-vaccine era London data, we also explored the impact of increasing or decreasing transmission intensity on genetic diversity. The t_MRCA_ reduces considerably as the transmission rate increases, as a result of increasingly severe population bottlenecks between outbreaks (Figure S2).

### Spatiotemporal Dynamics

#### Two populations

Perturbing population dynamics in one population can have an impact on neighboring communities. With even low levels of mixing, changes in genetic diversity in one population can be reflected in a neighboring community, despite the negligible effect on population size in the latter. Under intermediate vaccination coverage, the increase in genetic diversity in the targeted population can also be observed in the linked population (Figure 3) through importation and propagation of external lineages. With sufficiently high mixing, the genetic composition of both populations remains nearly identical across time.

**Figure 3.**
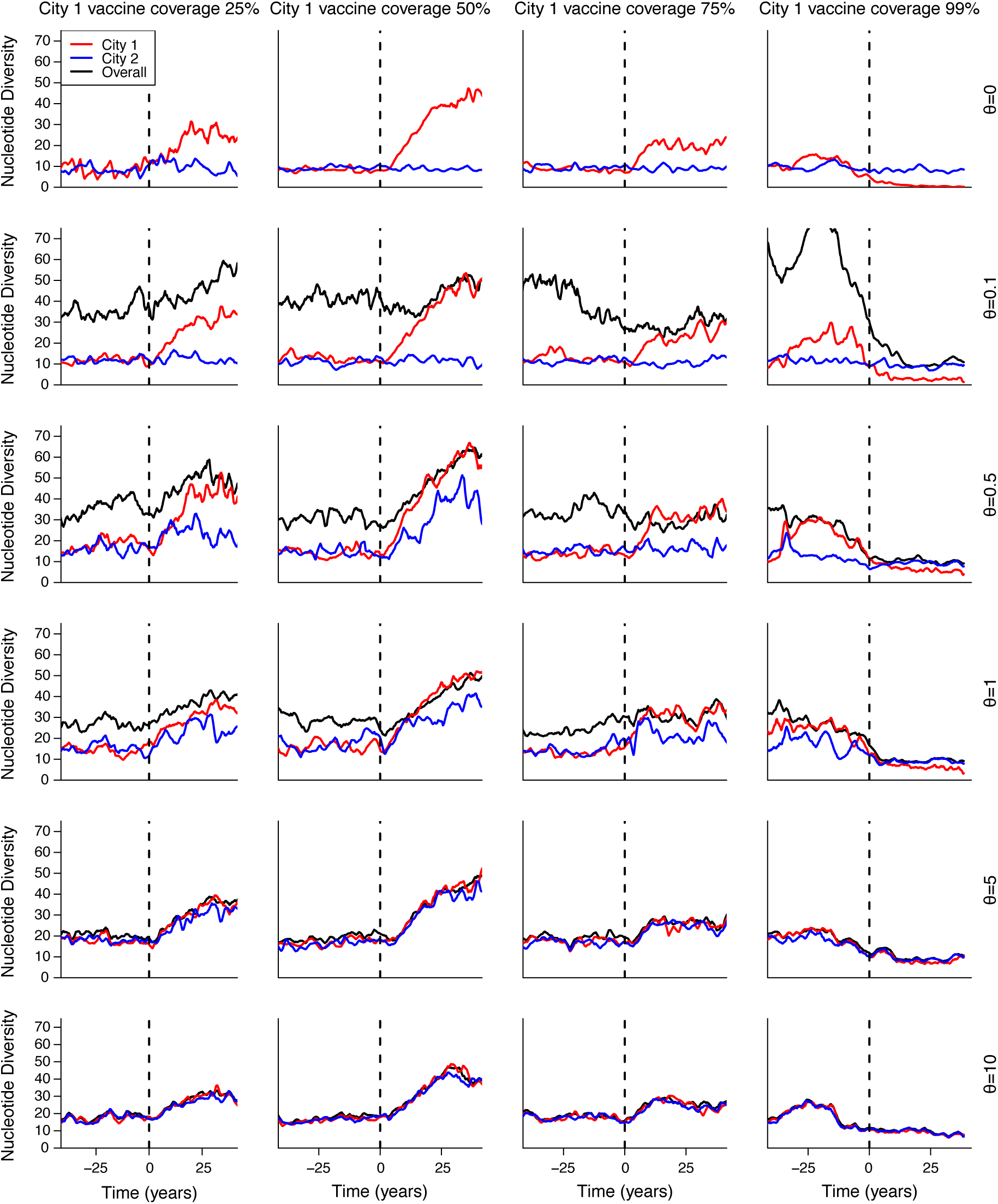
Vaccination in one population impacts neighboring communities. We simulated two equally sized populations with in phase biennial epidemics and a specified degree of mixing (theta), and introduced vaccination to one population, monitoring the sampled genetic diversity in each over time.

Populations experiencing out of phase epidemics were likely to experience a more rapid disruption of genetic composition. By seeding each population with its own distinct genotype, we could measure the length of their coexistence. Extinction of one strain was considerably more rapid when outbreaks were out of phase, such that a population with low between-epidemic incidence was under considerable external infectious pressure from the other population, experiencing a large outbreak (Figure 4).

**Figure 4.**
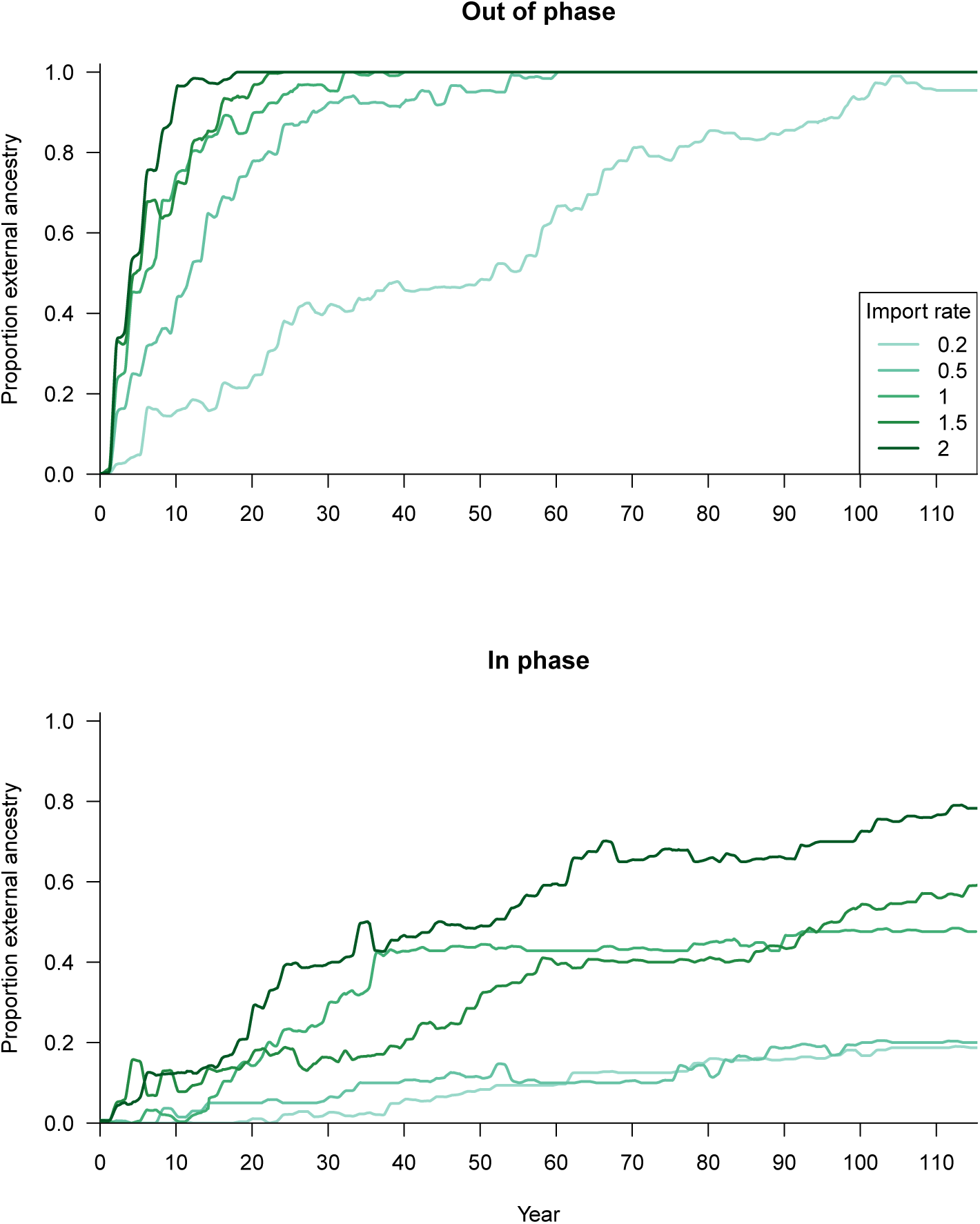
Strain replacement from external source. We simulate a major city with endemic monotypic measles. At time zero, cases may be imported from a second city with a second genotype circulating. For various rates of case importation, we measure the proportion of cases infected with the external genotype. We model the second city (A) out of phase, in which biennial outbreaks occur in alternate years in each city, and (B) in phase, in which biennial outbreaks occur simultaneously in each city. Importation rate is proportional to incidence in the external city, and is approximately the expected number of imported cases in each biweek at peak transmission. Average proportions across 25 simulations are shown.

#### Core-satellite metapopulation

We simulated populations comprising one major city and several minor cities, generating samples for 80 years pre- and post-vaccination (Figure S3). We found that after introducing vaccination at 50% coverage, measles remained endemic in the major city, and as previously, genetic diversity increased. In smaller cities (<200,000 inhabitants), the probability of extinction increased with vaccination introduction (Figure S4). However, the effect on expected genetic diversity was marginal (Figure 5). However, extinction became slightly rarer post-vaccination for larger cities (Figure S4). The stochastic loss and persistence of strains between seasons was the major driver of local fluctuations of genetic diversity. Global genetic diversity mirrored the dynamics within the endemic population centers, exhibiting an increase post-vaccination (Figure 5).

**Figure 5.**
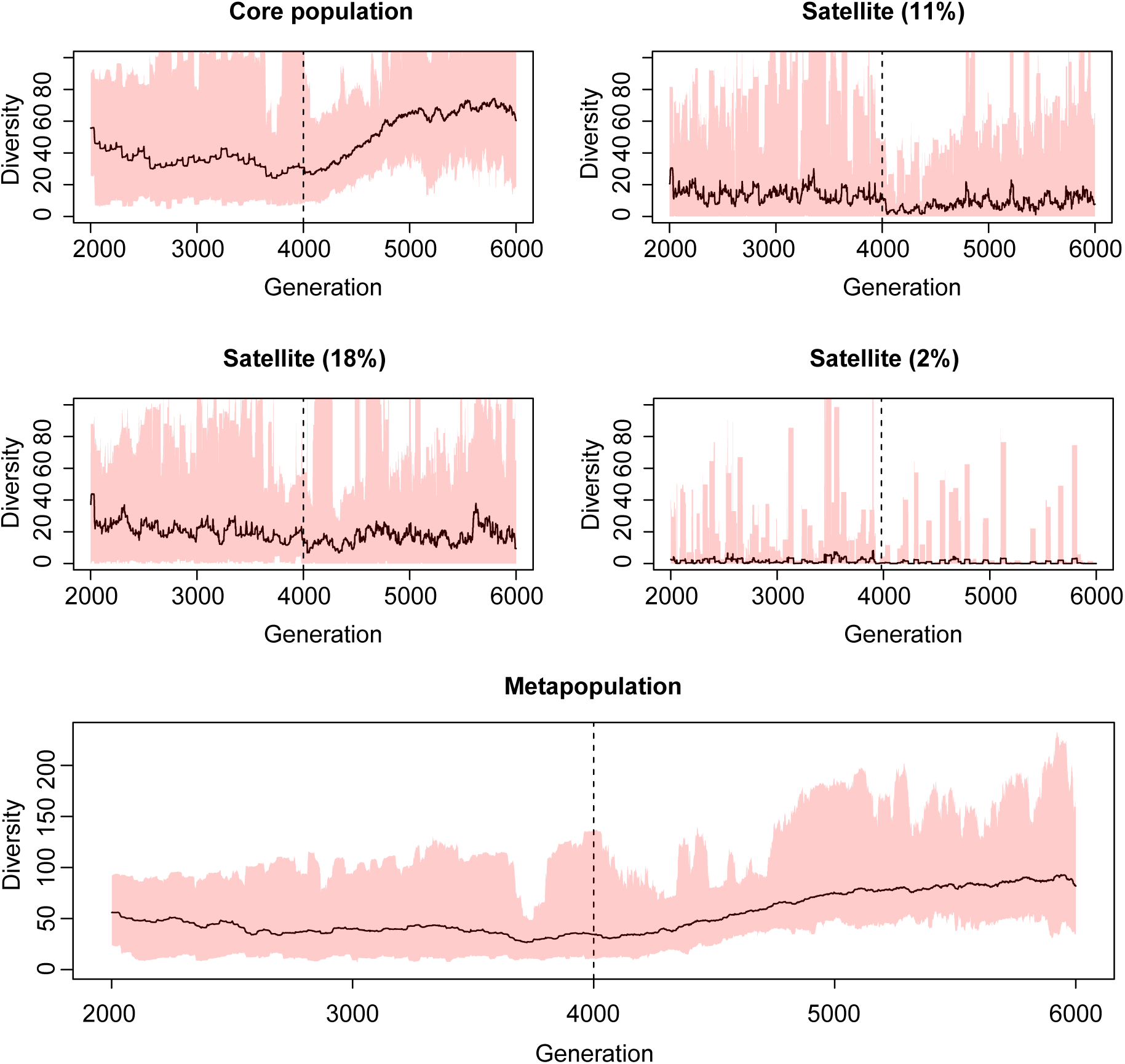
Mean genetic diversity across a metapopulation. The mean genetic diversity in a core-satellite metapopulation pre- and post-vaccination across 25 simulations. From generation 4000 (dashed line) 50% of the individuals in all populations were vaccinated. Shown are the core population (top left), and three example satellite populations, with populations expressed as a percentage of the core populations shown in parentheses. The bottom figure shows the overall metapopulation diversity. The shaded area represents the rage of diversity.

### GenBank sequence data

We considered the relationship between the monthly harmonic mean incidence and the observed genetic diversity for two countries in which measles remains endemic, China and India, as well as Europe (Figure 6; Table S1). While there were relatively few years with both incidence data and sequence data, we found that genetic diversity increased significantly with harmonic mean incidence for China (H1) (p=0.007) and Europe (B3) (p=0.02) as well as overall (p=0.003). Years with lower harmonic mean incidence appeared to have a greater variation in diversity, a result also observed in a simulated metapopulation setting (Figure S6). Once an endemic population becomes sufficiently small, the probability that an external strain (potentially a genetically distinct strain of the same genotype) can be imported and co-circulate becomes greater. This can lead to a higher diversity despite a lower incidence. Conversely, a disease-free community can be reseeded by a single strain, which can lead to a much lower diversity.

**Figure 6.**
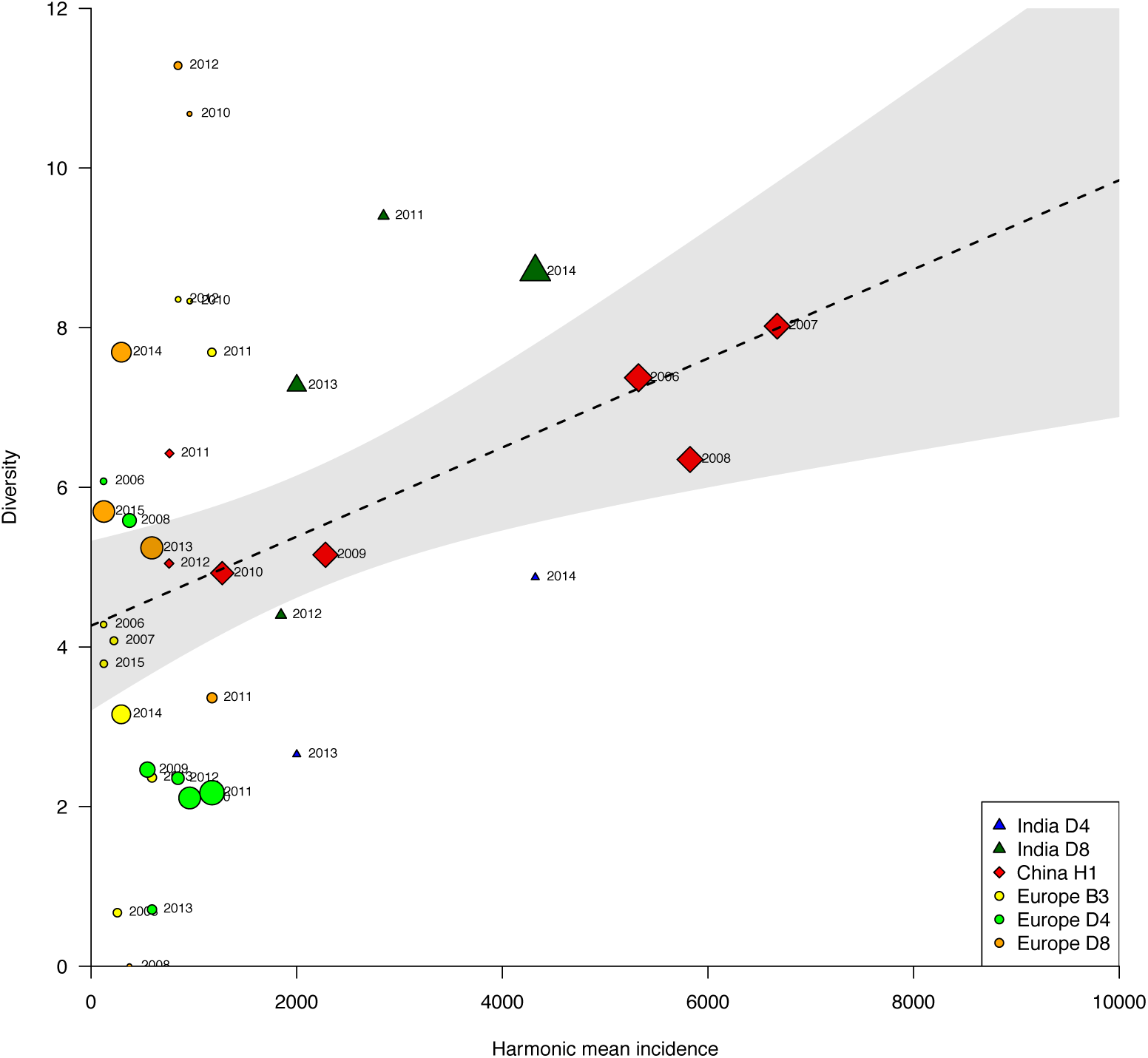
Nucleotide diversity increases with harmonic incidence. For India, China and Europe, we plot the harmonic mean measles incidence over the previous two years against the annual genetic diversity of the major genotypes circulating in each region. Point colors denote the country/genotype combination, while point size reflects the number of sequences available for each year. The dashed line denotes the weighted regression line while pooling all data points, with the 95% confidence interval shaded gray.

Measles incidence in China gradually reduced between 2005 and 2012, with both the annual peak and inter-epidemic bottleneck reducing, leading to a corresponding gradual reduction in genetic diversity of the endemic H1 genotype (Figure S5, top right). In contrast, genetic diversity of the entire measles collection increased, as introductions of different genotypes became more common (Figure S5, top left). Germany experienced measles outbreaks in 2011, 2013 and 2015, but genetic diversity in both the dominant genotype (D8) and across the whole population was greater in the intervening non-outbreak years (Figure S5, center). This may be due to extinction of the outbreak strain, followed by multiple sporadic introductions in the non-outbreak years. The USA experienced an outbreak in 2014, but its effect on observed genetic diversity both for the common genotype (B3) and the population overall was negligible (Figure S5, bottom). These data support our simulation findings that the relationship between genetic diversity and incidence patterns are far from straightforward, particularly in settings where measles is not endemic.

## Discussion

### Main results

Our findings suggest that a narrower population bottleneck between epidemics is expected to result in reduced pathogen genetic diversity, although considerable stochastic variation exists. Introducing vaccination will decrease the epidemic size but may increase the size of the population bottleneck between epidemics, leading to a larger expected genetic diversity. We found that at intermediate vaccination coverage levels (25-75%), a considerable increase in genetic diversity was expected, alongside increased variation around this value. While we have used measles dynamics as the basis of our simulations, we expect the bottleneck to play a key role in the genetic diversity observed in other seasonally-driven immunizing infectious disease systems too.

Building upon previous work (8), we have highlighted the challenges associated with the interpretation of genetic diversity in the context of disease dynamics for periodically fluctuating populations. Population bottlenecks, rather than peaks, characterize the expected levels of genetic diversity, although are associated with considerable short-term stochasticity. While introducing vaccination reduces incidence, it can allow increased transmission between major epidemics at intermediate coverage levels. Under the commonly used seasonally-forced SIR model, the rate of transmission between epidemics is dependent on the size of the susceptible population remaining post-outbreak, which is expected to be more heavily depleted after an outbreak with a higher force of infection, whether through increased R_0_, or a larger initial susceptible population. Reducing the growth rate of the susceptible population (i.e. reducing birth rate or increasing vaccination coverage) leads to a smaller susceptible population at the start of a season, and the depletion of susceptibles during the outbreak is much slower. This results in a greater number of susceptibles remaining once the outbreak is over, with reduced seasonal transmission rates preventing outbreaks becoming drawn out over time. This work also highlights the important role of stochasticity in the between-epidemic population dynamics driving observed genetic diversity in the following season. As such, care should be taken interpreting an observed increase or decrease in genetic diversity from one season to the next. It was previously demonstrated that genotype switching could frequently be ascribed to stochastic effects, rather than vaccination coverage (19). Our study demonstrates that similar principles apply to fluctuations in within-genotype diversity.

We were able to explore the relationship between genetic diversity and harmonic mean incidence in India, China and Europe using sequence data from GenBank. While the scope of this study was somewhat limited by the availability of monthly incidence data and/or sufficient numbers of sampled sequences, we observed a significantly positive trend between harmonic mean incidence and genetic diversity. There was a greater variation in diversity for smaller values of harmonic mean incidence due to the increased potential for co-circulating imported strains. Since sequences on GenBank originate from a variety of studies and surveillance frameworks, it is unlikely that these represent a truly random sample from the populations in question. With a larger dataset, it may be possible to investigate more complex models of observed diversity, accounting for year-to-year dependencies and sample size. Furthermore, most of the sequencing conducted for measles has focused only on the N-450 sequence. Whole genome sequencing is likely to provide greater resolution (40); in practice several mutations were observed during a single outbreak in Canada in 2010 (41). WHO is developing recommendations for the use of whole genome sequencing for tracing transmission of measles virus (42) and would allow changes in genetic diversity to be measured on a finer time scale.

### Previous measurements of measles genetic diversity

Previous studies have linked a change in genetic diversity of measles to the impact intervention measures. Kremer et al. suggested that a decline in the genetic diversity of the D6 genotype in Europe in 2005-06 relative to the 1990s and early 2000s may be ascribed to enhanced vaccination (13). Since vaccination coverage in Europe was already high on average in the 1990s (43), such a finding may indeed be consistent with our theoretical findings, as we also observed a reduction in diversity when increasing vaccination beyond 50% coverage. However, we found that diversity across a metapopulation was driven by large subpopulations with endemic disease, so an observed reduction in diversity could also potentially result from deeper bottlenecks in such regions, arising either by chance, or indeed by large preceding outbreaks. Another study suggested the greater diversity of measles in Nigeria compared to the Democratic Republic of Congo (DRC) in the mid-2000s may be due to the higher vaccine coverage in the latter (24). As the study acknowledges, differences in demographics and movement patterns in the two countries makes comparisons challenging. We have furthermore shown that changes in diversity may also arise through stochastic variation or the influence of linked and potentially unobserved external populations.

Measles in China remains endemic, despite ongoing efforts to increase vaccination coverage. The genotype H1 has been dominant for many years (44), suggesting that incidence remains sufficiently high year round for the probability of strain replacement by imported cases to be small. The reduction in diversity of H1 genotypes in China between 2006 and 2007 was linked to the measles elimination action plan put in place by the Chinese Ministry of Health in 2006 (23). However, similar monthly incidence reports were observed in each year, and our theoretical findings suggest that such season-to-season fluctuations in diversity are frequently observed by chance. Similarly, a reduction in diversity between 2009 and 2010 may not necessarily be a direct effect of a nationwide supplementary immunization activity in 2010 as has been suggested (22).

### Future work

Genetic surveillance is an important tool in investigating transmission patterns of communicable diseases (45), including measles (46). The ability to discriminate between endemic and imported cases is crucial in determining appropriate control measures to reduce transmission (15, 47). Further work is still required to fully understand the relationship between epidemiological and evolutionary dynamics. While genetic diversity is commonly used as a metric to measure the success of measles reduction, it is possible to use more sophisticated approaches to infer explore population dynamics using sequence data. Stack et al. demonstrated that with a carefully designed sampling strategy, it is possible to generate skyline plots which broadly capture seasonal dynamics using the software BEAST (8). However, the study did not investigate the impact of vaccination, movement between multiple population centers, nor the sensitivity of the approach to detect differences in peak incidence or trough depth. Furthermore, the study highlighted the sensitivity of the approach to different sampling strategies. Later work demonstrated that a standard Bayesian skyline approach failed to identify seasonal dynamics, though with a properly specified epidemiological model, these dynamics can be recaptured (48, 49). Further research is needed to determine whether such phylogenetic analyses in combination with epidemic models could shed more light on the impact of infection control measures for seasonal outbreaks, allowing one to correctly infer the changes in peak incidence rather than bottleneck severity.

While our study focused on measles dynamics, other seasonally-driven epidemics may have differing characteristics. While we believe that the inter-epidemic bottleneck will be important in almost all contexts, larger outbreaks with a higher rate of mutation (such as influenza), may generate sufficient within-season diversity to diminish the role of pre-outbreak diversity. Our study is also limited in assuming that diversity arises purely through genetic drift; while this is appropriate for measles, recombination and selection pressure will play an important role for many other pathogens. Furthermore, we assumed that all strains were identical in terms of virulence and vaccine efficacy. Further work is required to explore the impact of these factors on pathogen population diversity.

## Acknowledgements

This work was supported by the Bill & Melinda Gates Foundation, the US Centers for Disease Control and the RAPIDD program of the Science and Technology Directorate, Department of Homeland Security and the Fogarty Center, US National Institutes of Health.

## Supplementary Figures

**Figure S1.**
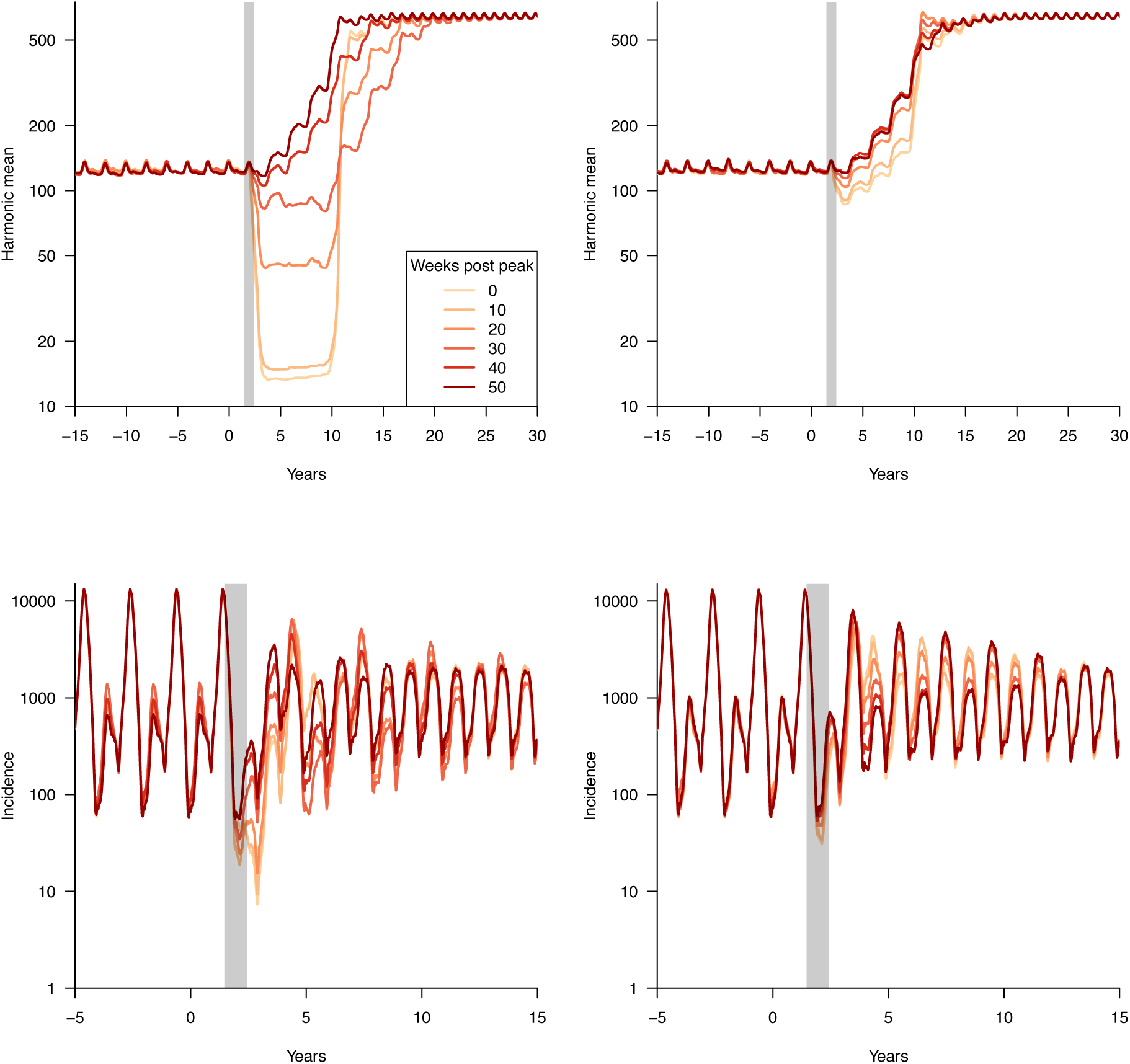
Timing and speed of vaccine introduction affects short term dynamics. (A) The effect of instantaneous introduction of 50% vaccination coverage on the harmonic mean incidence (10 year sliding window). Vaccination was introduced at different time points following a major epidemic; more rapid vaccination leads to a deeper bottleneck, and greater probability of extinction. Incidence is shown in panel (C). (B) Gradual vaccination introduction. When introducing vaccination, 20% of the population is covered, and coverage levels increase linearly to 50% over 10 years. Incidence is shown in panel (D).

**Figure S2.**
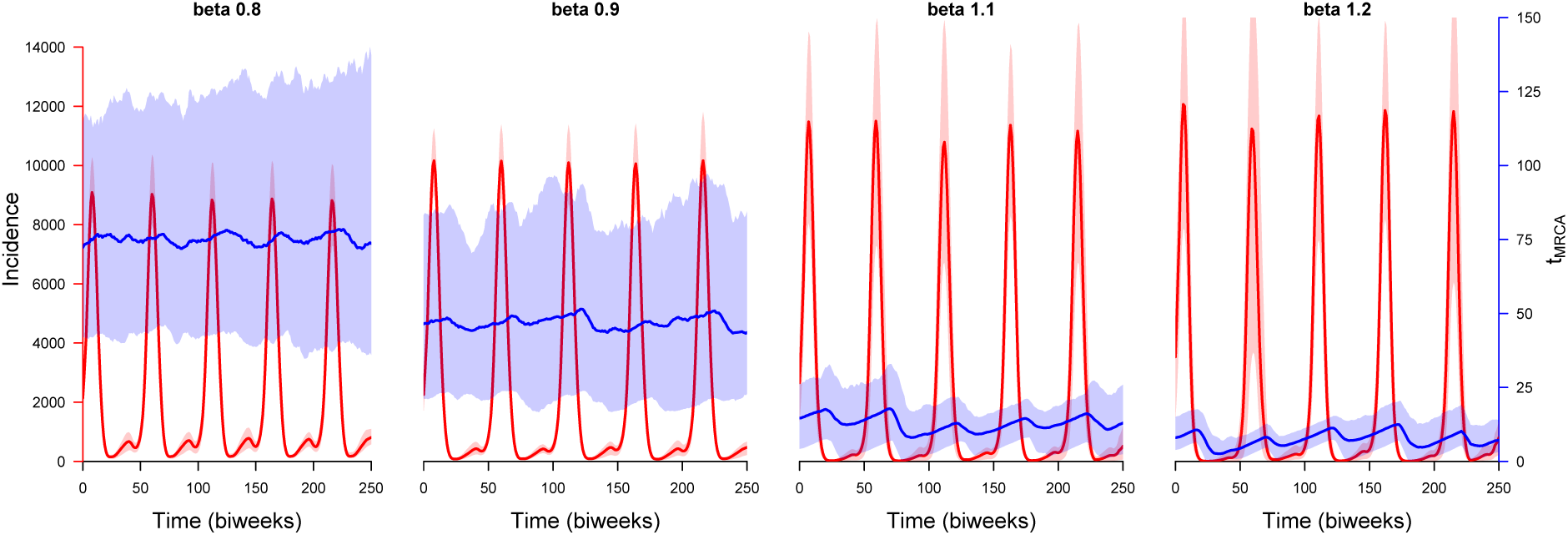
Simulated measles incidence (red) and mean time to most recent common ancestor (t_MRCA_) (blue). We varied the transmission rate (uniformly multiplying the fluctuating seasonal transmission rates by a factor beta) and plotted the average incidence and t_MRCA_ over a period of ten years, averaging both across 50 simulations. Shaded areas represent the interquartile range.

**Figure S3.**
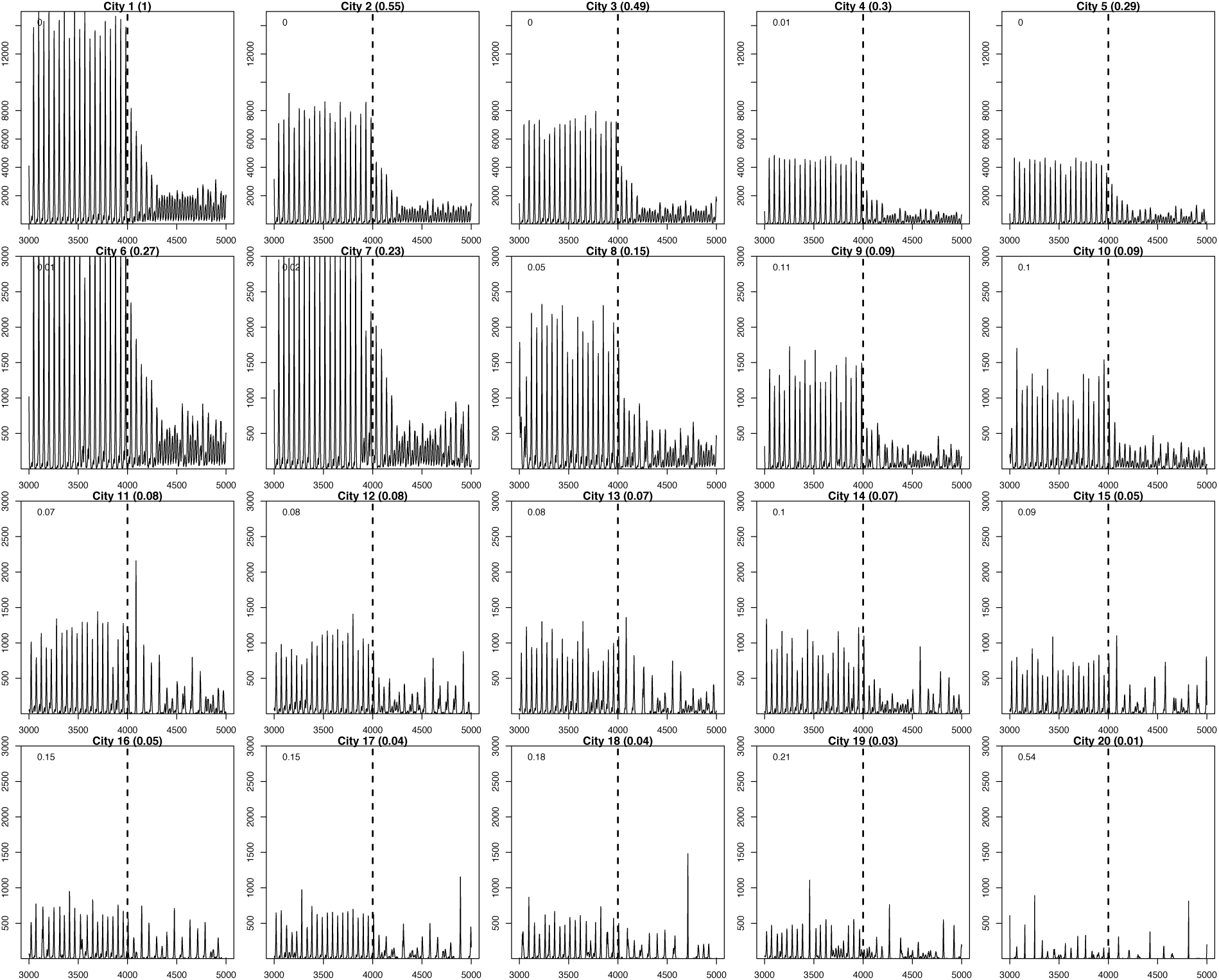
Pre- and post-vaccination incidence in a 20 city metapopulation. Vaccination is introduced at generation 4000 at 20% coverage, gradually increasing to 50% within 10 years (260 generations). Cities are shown in decreasing order of size (given relative to the major city in parentheses in plot titles). Values in the top left of each plot give the proportion of generations pre-vaccine for which incidence in that city is zero.

**Figure S4.**
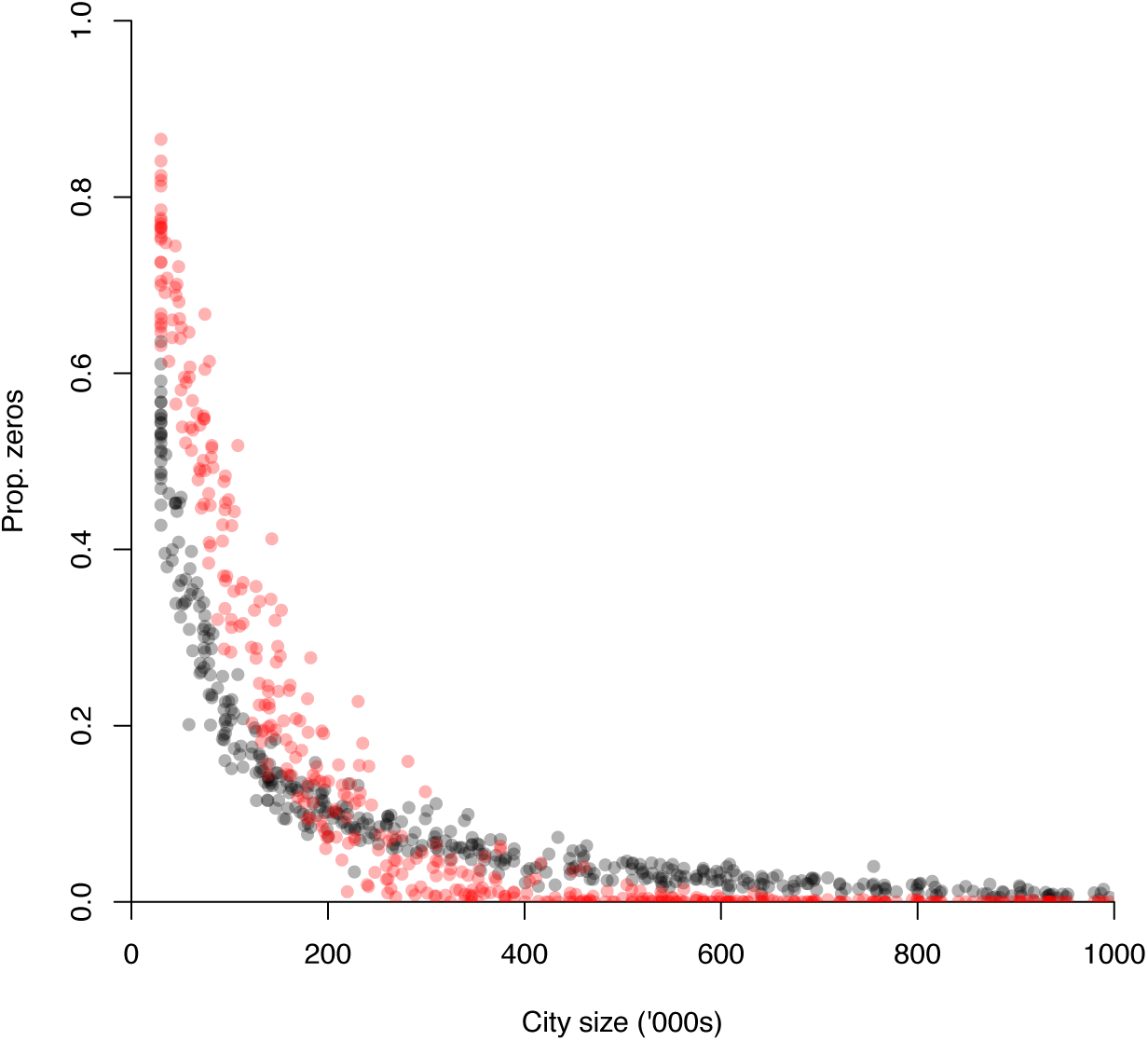
Frequency of extinction by city size. The proportion of biweeks in which incidence is zero in cities of various size in a 20 city metapopulation. Proportion of zeroes both pre-vaccination (black) and post-vaccination (red) are shown. Vaccination coverage was 50%.

**Figure S5.**
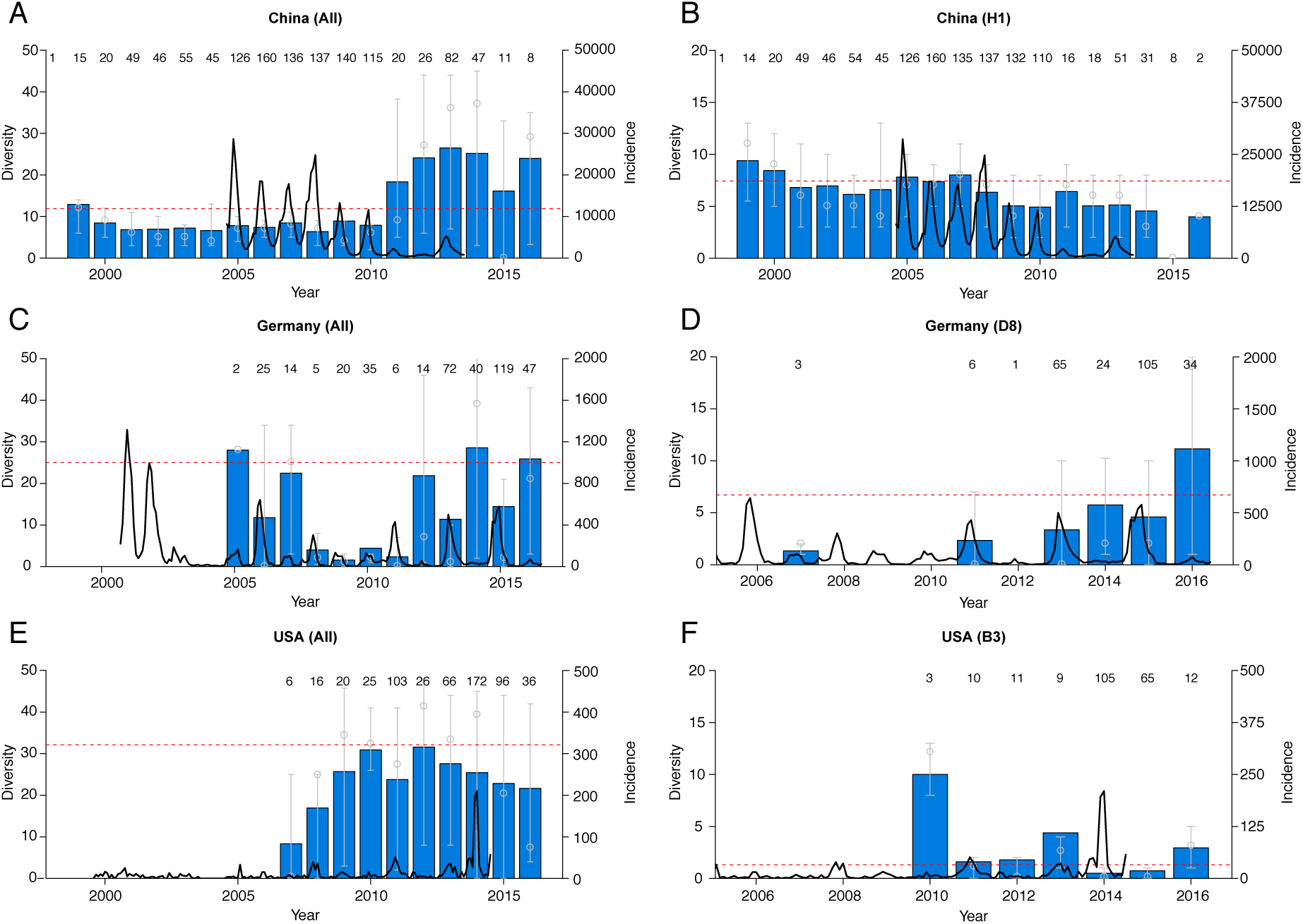
Observed measles genetic diversity by country and genotype. We obtained all publicly available measles sequence data (covering at least N-450) for China (top), Germany (center) and the USA (bottom) from the GenBank database (retrieved October 2017). We show the overall diversity, as well as that of the most common genotype. Annual mean nucleotide diversity is given by the colored bars; median and interquartile ranges are represented by the open circles and vertical lines. Annual numbers of sequences are given above each bar. The overall diversity across time is shown by the dashed red line. Monthly incidence is given by the black line.

**Figure S6.**
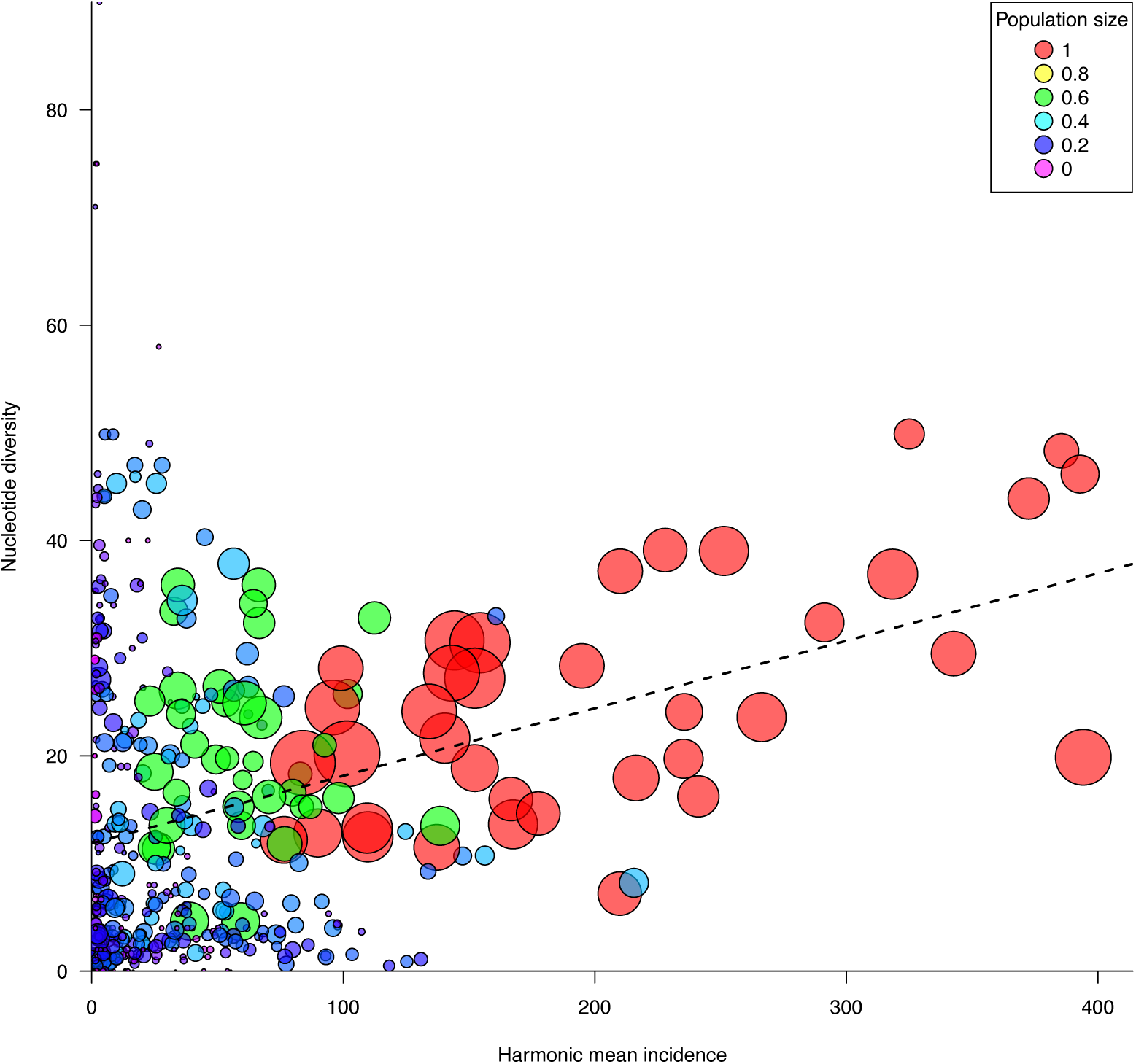
Nucleotide diversity in simulated populations increases with harmonic mean incidence. Using simulated data from a core-satellite metapopulation, for each year we plot the harmonic mean incidence over the past two years against the sampled nucleotide diversity, analogous to Figure 6. Point colors represent the relative size of the sampled populations; a relative size of 1 corresponds to the core population of size 3 million. Point sizes denote the number of samples from each year. The dashed line denotes the weighted regression line while pooling all data points.

**Table S1.** Breakdown of NCBI Genbank measles samples by country and genotype.

| Region | Total samples | Main genotypes |
| --- | --- | --- |
| Europe | 1851 | D8 (622); D4 (610); B3 (412) |
| China | 1007 | H1 (925) |
| India | 1044 | D8 (797); D4 (165) |
| USA | 587 | B3 (229); D8 (156) |
| Germany | 409 | D8 (253) |

